# Seeded aggregation of ANXA11 induces prion-like propagation, TDP-43 co-pathology and nucleocytoplasmic transport defects

**DOI:** 10.64898/2026.03.24.713853

**Authors:** Haiyang Luo, Honglin Zheng, Yongting Lu, Chenyang Liu, Kang Zhang, Suying Duan, Hang Zhang, Yaochong Zhang, Yaxuan Song, Tuo Wang, Han Liu, Na Zhang, Zongping Xia, Yuming Xu

**Affiliations:** Department of Neurology, First Affiliated Hospital of Zhengzhou University, Zhengzhou University, Zhengzhou, Henan, China; Department of Neurology, Beijing Tiantan Hospital, Capital Medical University; Beijing, China; Department of Pathology, Second Affiliated Hospital of Zhengzhou University, Zhengzhou University, Zhengzhou, Henan, China; Clinical Systems Biology Laboratories, Translational Medicine Center, First Affiliated Hospital of Zhengzhou University, Zhengzhou, Henan, China; NHC Key Laboratory of Prevention and Treatment of Cerebrovascular Disease, Zhengzhou, Henan, China; Henan Key Laboratory of Cerebrovascular Diseases, Zhengzhou University, Zhengzhou, Henan, China; Institute of Neuroscience, Zhengzhou University, Zhengzhou, Henan, China

## Abstract

Mutations in ANXA11 cause amyotrophic lateral sclerosis (ALS) and frontotemporal dementia (FTD), yet the mechanisms linking ANXA11 dysfunction to neurodegeneration remain poorly defined. Recent cryo-EM studies revealed heteromeric ANXA11-TDP-43 filaments in patient brains, suggesting a direct pathological connection between these two ALS-associated proteins. However, whether ANXA11 possesses intrinsic amyloidogenic properties and how its aggregation relates to TDP-43 proteinopathy remain unknown. Here, we demonstrate that ANXA11 undergoes liquid-liquid phase separation and subsequently matures into amyloid fibrils through a liquid-to-solid phase transition. ANXA11 fibrils exhibit prion-like properties, including self-templating seeding activity and intercellular propagation in human iPSC-derived neurons. Strikingly, ANXA11 fibrils induces pathological conversion of TDP-43, including hyperphosphorylation, accumulation in detergent-insoluble fractions, and formation of cytoplasmic aggregates. TurboID proximity-labeling proteomics further revealed aggregation-dependent enrichment of nuclear pore complex and nucleocytoplasmic transport factors in the ANXA11 aggregate-proximal proteome. Consistently, ANXA11 aggregation was associated with nuclear envelope abnormalities, altered nucleoporin distribution, impaired mRNA export, and progressive neuronal toxicity in iPSC-derived neurons. Together, these findings establish ANXA11 as an intrinsically amyloidogenic, phase-transition-competent protein whose seeded assemblies propagate between cells, induce TDP-43 co-pathology, and are linked to nucleocytoplasmic transport defects and neuronal injury, thereby providing a mechanistic framework for ANXA11-associated ALS/FTD pathogenesis.

**Graphic Abstract:** 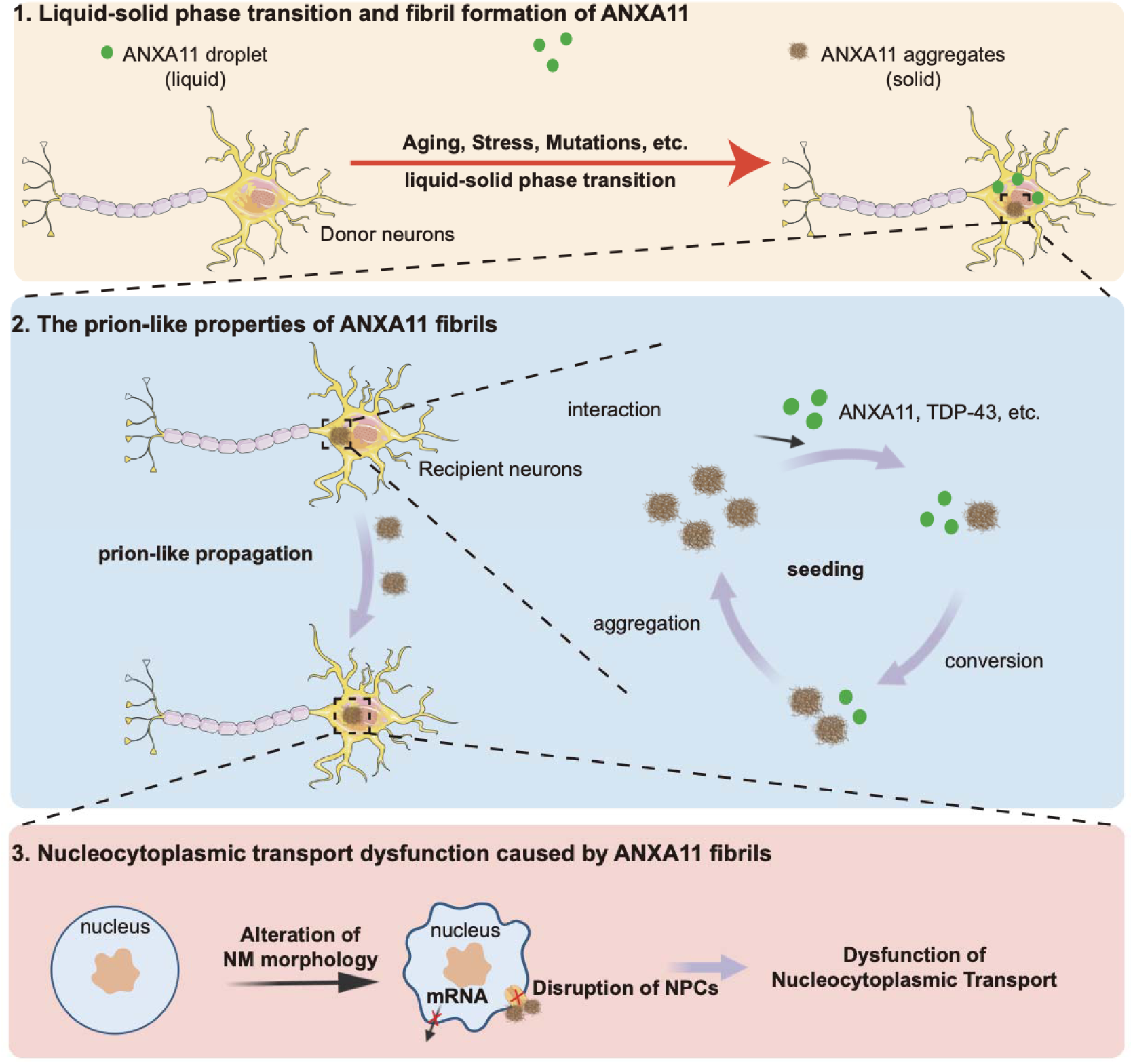

**Highlights:**

1. ANXA11 undergoes liquid-liquid phase separation and matures into amyloid fibrils through a liquid-to-solid phase transition.
2. ANXA11 fibrils supports homotypic seeding and propagate within SH-SY5Y cells and iPSC-derived neurons.
3. ANXA11 fibrils induce TDP-43 pathological conversion, including phosphorylation and accumulation in insoluble aggregates.
4. ANXA11 aggregation is associated with nuclear pore complex remodeling, impaired mRNA export, and neuronal toxicity.

**In Brief:** ANXA11 undergoes phase separation and matures into prion-like amyloid fibrils through a liquid-to-solid transition. These seeded assemblies propagate between cells and human neurons, induce TDP-43 pathological conversion, and are associated with remodeling of nuclear pore complexes, impaired mRNA export, and progressive neuronal toxicity.

ANXA11 forms seeded amyloid assemblies that spread between cells, induce TDP-43 pathology, and disrupt nucleocytoplasmic transport.

## INTRODUCTION

Cytoplasmic aggregation and nuclear clearance of TAR DNA-binding protein 43 (TDP-43) are key pathological hallmarks of amyotrophic lateral sclerosis (ALS) and frontotemporal dementia (FTD), affecting ∼97% and ∼45% of cases, respectively (Neumann et al., 2006; Arai et al., 2006; Ling et al., 2013). Recent cryo-electron microscopy (cryo-EM) studies have expanded the repertoire of proteins implicated in TDP-43 proteinopathies, revealing unexpected complexity in amyloid filament composition. Notably, Annexin A11 (ANXA11) was identified as a component of heteromeric amyloid filaments together with TDP-43 in frontotemporal lobar degeneration with TDP-43 pathology type C (FTLD-TDP-C) (Arseni et al., 2024). Additional immunohistochemical analyses confirmed ANXA11-positive inclusions across several other TDP-43 proteinopathies. These observations raise the possibility that ANXA11 is not merely associated with pathological inclusions, but may actively contribute to amyloid assembly and downstream cellular dysfunction. However, whether ANXA11 possesses intrinsic fibrillization capacity and how its aggregation perturbs cellular functions remain unresolved.

ANXA11 is a Ca²□-dependent phospholipid- and RNA granule-binding protein containing a long N-terminal intrinsically disordered region (IDR), a domain architecture shared by numerous prion-like proteins that undergo liquid-liquid phase separation (LLPS) to form dynamic biomolecular condensates (Molliex et al., 2015; Patel et al., 2015; Nahm et al., 2020). Dominant mutations in ANXA11 cause familial ALS (Smith et al., 2017; Jiang et al., 2022), establishing a direct genetic link between ANXA11 dysfunction and neurodegeneration. Increasing evidence suggests that LLPS-competent proteins can transition from liquid-like condensates to solid amyloid assemblies under stress or aging conditions (Murakami et al., 2015; Wegmann et al., 2018), This phase-to-aggregate continuum provides a plausible mechanistic framework by which ANXA11 could shift from a physiological condensate-associated state to a pathological assembly state. Nevertheless, the capacity of ANXA11 to undergo such a liquid-to-solid transition and the consequences of such transition have not been systematically defined.

Numerous aggregation-prone proteins, including α-synuclein, tau, TDP-43, hnRNPA1, and FUS, can self-assemble into β-sheet-rich fibrils and propagate this conformational conversion in a prion-like manner, thereby amplifying misfolded species across cells (Polymenidou and Cleveland, 2011; Nonaka et al., 2013). Prion-like conversion has thus emerged as a unifying mechanism underlying the spread of protein aggregation within the nervous system. The heteromeric ANXA11-TDP-43 filaments described above raises the possibility that ANXA11 assemblies may undergo homotypic self-templating potentially cross-seed TDP-43 aggregation, a mechanism that could explain the co-pathology observed in patients; whether such templating events indeed occur remains to be determined. Beyond assembly mechanisms, the cellular pathways most directly affected by ANXA11 aggregation remain poorly understood. In ALS/FTD, disruption of nucleocytoplasmic transport and nuclear pore complex (NPC) integrity are recurrent pathological themes (Zhang et al., 2015; Chou et al., 2018), and TDP-43 itself has been implicated in NPC perturbation. Whether ANXA11 assemblies compromise this pathway and contribute to neuronal toxicity have yet to be directly tested.

Here, we investigate the molecular behavior and pathological consequences of ANXA11 aggregation. We show that ANXA11 undergoes liquid-liquid phase separation and transitions into amyloid-like fibrils with self-templating seeding activity that propagates between cells and human iPSC-derived neurons. These fibrils trigger pathological TDP-43 conversion and proteomic reorganization involving nucleocytoplasmic transport factors, leading to impaired mRNA export and neuronal toxicity. Together, our findings define a mechanistic link between ANXA11 phase transition, seeded propagation, TDP-43 co-pathology, and transport dysfunction in ALS/FTD-related neurodegeneration.

## Results

### ANXA11 forms amyloid fibrils with self-templating seeding activity

ANXA11 contains an extensive N-terminal low complexity domain (LCD), as predicted by PONDR and IUPred2A analyses (Figure 1A). Given that IDR-containing proteins frequently undergo liquid-liquid phase separation (LLPS), we assessed the phase behavior of recombinant full-length ANXA11 in vitro. Under macromolecular crowding conditions (10% PEG8000), ANXA11 solutions became turbid in a concentration-dependent manner, whereas bovine serum albumin (BSA) control did not exhibit phase separation (Figure 1B). Alexa Fluor 488-labeled ANXA11 formed micron-scale spherical droplets visible by fluorescence and differential interference contrast (DIC) microscopy (Figure 1C). Fluorescence recovery after photobleaching (FRAP) revealed that freshly formed ANXA11 droplets are highly dynamic, with rapid fluorescence recovery characteristic of liquid-like condensates (Figure 1D-E).

**Figure 1.**
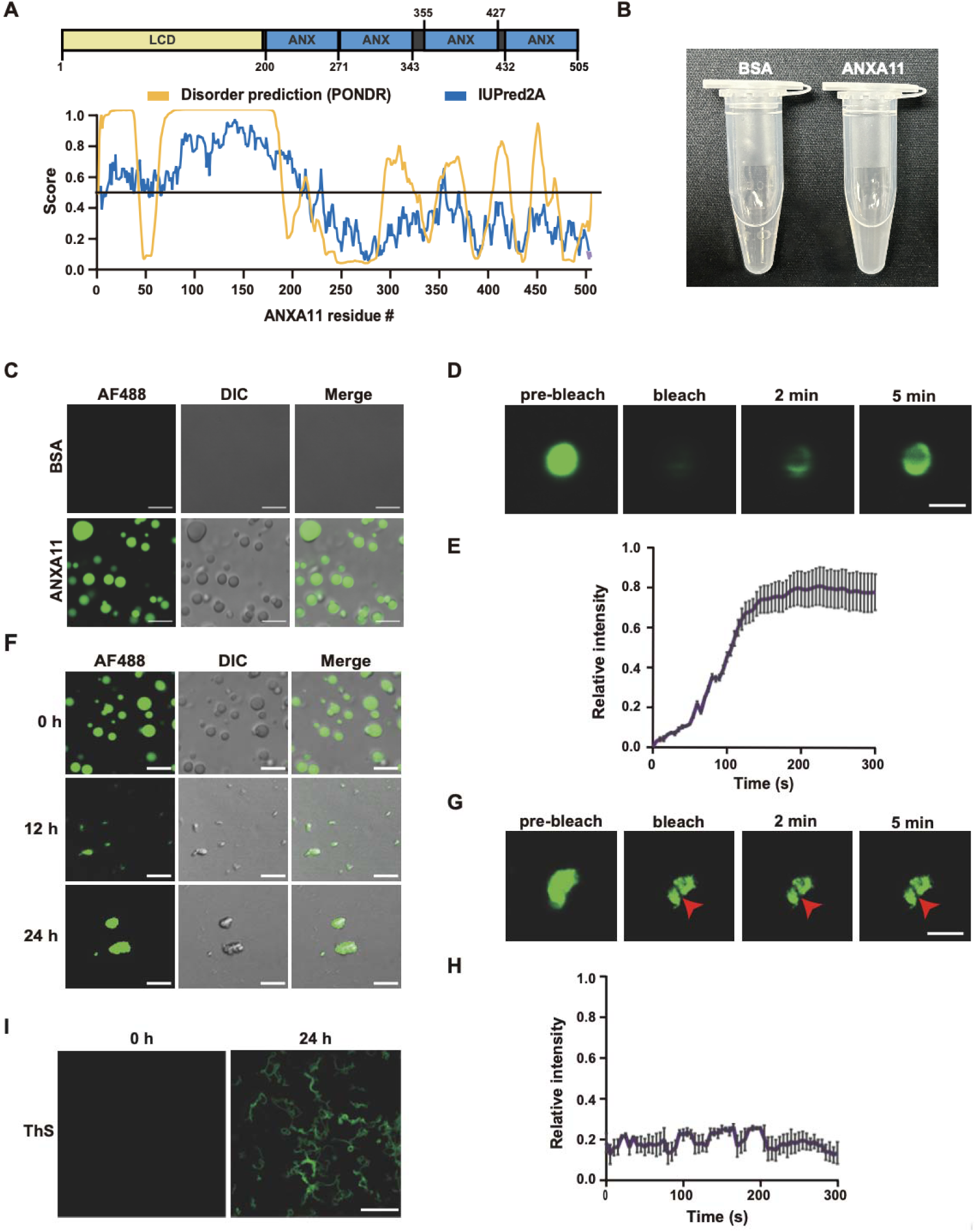
ANXA11 undergoes LLPS and liquid-to-solid phase transition in vitro. (A) Domain architecture of ANXA11 showing the N-terminal low complexity domain (LCD) and C-terminal annexin repeats. Disorder prediction scores from PONDR (blue) and IUPred2A (orange) algorithms are shown. (B) Turbidity assay measuring macroscopic phase separation of recombinant ANXA11 solutions in vitro; BSA is a negative control. Data are representative of three independent experiments. (C) Representative fluorescence and differential interference contrast (DIC) images of AF488-labeled ANXA11 droplets formed under crowding conditions. Scale bar: 10 μm. (D) Fluorescence recovery after photobleaching (FRAP) analysis of freshly formed ANXA11 droplets. Representative images before bleaching, immediately after bleaching, and during recovery are shown. Red circle indicates bleached region. Scale bar: 5 μm. (E) Quantification of FRAP recovery kinetics for fresh ANXA11 droplets demonstrating liquid-like behavior shown in (D). Data represent mean ± SD from at least 10 droplets per condition from 3 independent experiments. (F) Time-series imaging of ANXA11 droplets during aging (0 h, 12 h, 24 h) showing progressive changes in morphology. Scale bar: 10 μm. (G) FRAP analysis of aged (24 h) ANXA11 aggregates. Representative images showing minimal fluorescence recovery after photobleaching. Scale bar: 5 μm. (H) Quantification of FRAP recovery kinetics for aged ANXA11 aggregates shown in (H) demonstrating solid-like behavior with negligible fluorescence recovery. Data represent mean ± SD from at least 10 aggregates per condition from 3 independent experiments. (I) Thioflavin S (ThS) staining of ANXA11 assemblies at 0 h versus 24 h, demonstrating acquisition of amyloid-like properties upon aging. Scale bar: 20 μm.

Liquid condensates formed by LCD-containing proteins can mature into more stable, solid-like assemblies over time—a process termed aging. To determine whether ANXA11 droplets undergo such liquid-to-solid phase transition, we incubated them at 37°C and monitored their biophysical properties over time. Time-series imaging revealed progressive changes in droplet morphology during aging (Figure 1F). FRAP analysis of aged ANXA11 assemblies showed minimal fluorescence recovery (Figure 1G-H), indicating conversion to a solid-like state. Thioflavin S (ThS) staining demonstrated that aged ANXA11 assemblies acquired amyloid-like properties, with strong ThS fluorescence at 24 hours compared to freshly formed droplets (Figure 1I). Together, these data show that ANXA11 fibrils possess robust self-templating seeding activity, a defining property of prion-like protein assemblies.

### Intracellular ANXA11 droplets mature into amyloid-like aggregates

To determine whether ANXA11 phase transitions occur in living cells, we expressed GFP-tagged ANXA11 in U2OS cells and monitored condensate dynamics over time. At early time points, ANXA11-GFP formed cytoplasmic puncta that exhibited liquid-like properties: FRAP analysis revealed rapid fluorescence recovery (Figure 2A-B), and time-lapse imaging captured fusion and fission events characteristic of liquid droplets (Figure 2C). These observations confirm that ANXA11 undergoes LLPS in the cellular environment.

**Figure 2.**
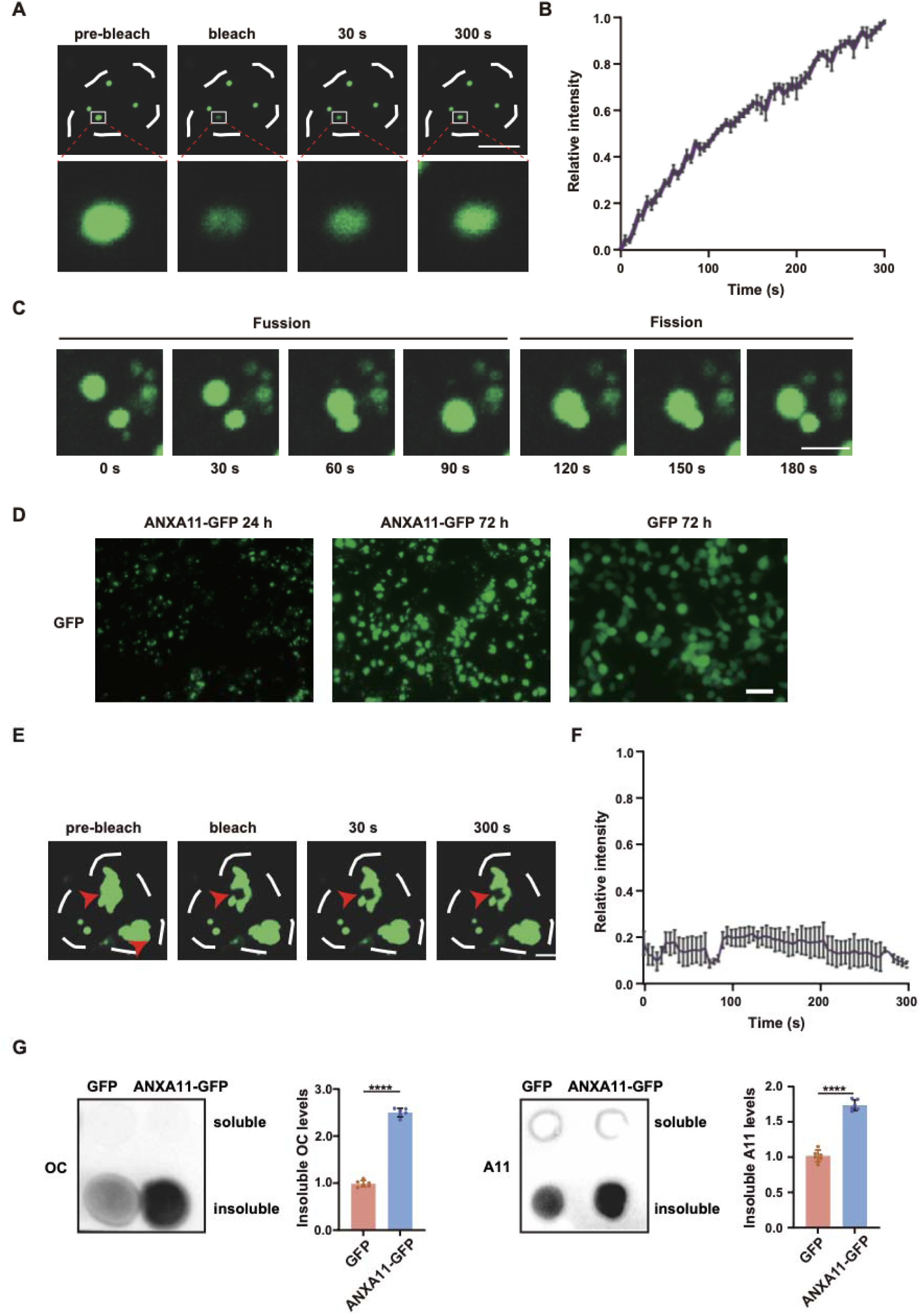
Intracellular ANXA11 condensates mature into amyloid-like assemblies. (A) FRAP analysis of nascent ANXA11-GFP droplets in U2OS cells. Representative images before bleaching, immediately after bleaching, and during recovery are shown. Scale bar: 5 μm. (B) Quantification of FRAP recovery kinetics for nascent intracellular ANXA11-GFP condensates shown in (A) demonstrating liquid-like behavior. Data represent mean ± SD from at least 10 condensates from 3 independent experiments. (C) Time-lapse imaging capturing fusion and fission dynamics of ANXA11-GFP condensates, confirming liquid-like properties. Time stamps are indicated. Scale bar: 5 μm. (D) Time-dependent aggregation of ANXA11-GFP showing morphological changes between 24 h and 72 h post-transfection. Scale bar: 10 μm. (E) FRAP analysis of 72 h-old ANXA11-GFP aggregates. Representative images showing minimal fluorescence recovery after photobleaching. Scale bar: 5 μm. (F) Quantification of FRAP recovery kinetics for 72 h-old ANXA11-GFP aggregates shown in (E) demonstrating solid-like behavior. Data represent mean ± SD from at least 10 aggregates from 3 independent experiments. (G) Dot blot analysis of cell lysates from ANXA11-GFP-expressing cells probed with conformation-specific antibodies OC (fibrillar amyloid) and A11 (prefibrillar oligomers); total protein stain serves as a loading control.

However, with prolonged expression, ANXA11 condensates underwent a striking transformation. Time-dependent imaging revealed progressive changes in condensate morphology between 24 and 72 hours post-transfection (Figure 2D). FRAP analysis of 72-hour-old ANXA11 assemblies showed minimal fluorescence recovery (Figure 2E-F), indicating that the condensates had converted to a solid-like state. To assess whether these solid-like inclusions acquired amyloid properties, we performed dot blot analysis using conformation-specific antibodies. Lysates from cells harboring aged ANXA11 inclusions showed strong reactivity with the OC antibody, which recognizes fibrillar amyloid structure, and the A11 antibody, which detects prefibrillar oligomers (Figure 2G). These findings demonstrate that intracellular ANXA11 condensates undergo a time-dependent liquid-to-solid phase transition and acquire amyloid-like properties, recapitulating the in vitro observations within a cellular context.

### ANXA11 forms amyloid fibrils with prion-like self-templating activity

To further characterize the amyloidogenic properties of ANXA11, we employed computational and biochemical approaches. FoldAmyloid prediction identified aggregation-prone regions within the ANXA11 sequence (Figure 3A). Congo red and Thioflavin S staining of aged ANXA11 assemblies confirmed their amyloid nature, with characteristic birefringence under polarized light and strong ThS fluorescence (Figure 3B). Transmission electron microscopy (TEM) revealed that while monomeric ANXA11 appeared as dispersed particles, aged ANXA11 formed fibrillar structures with characteristic amyloid morphology, including unbranched filaments (Figure 3C).

**Figure 3.**
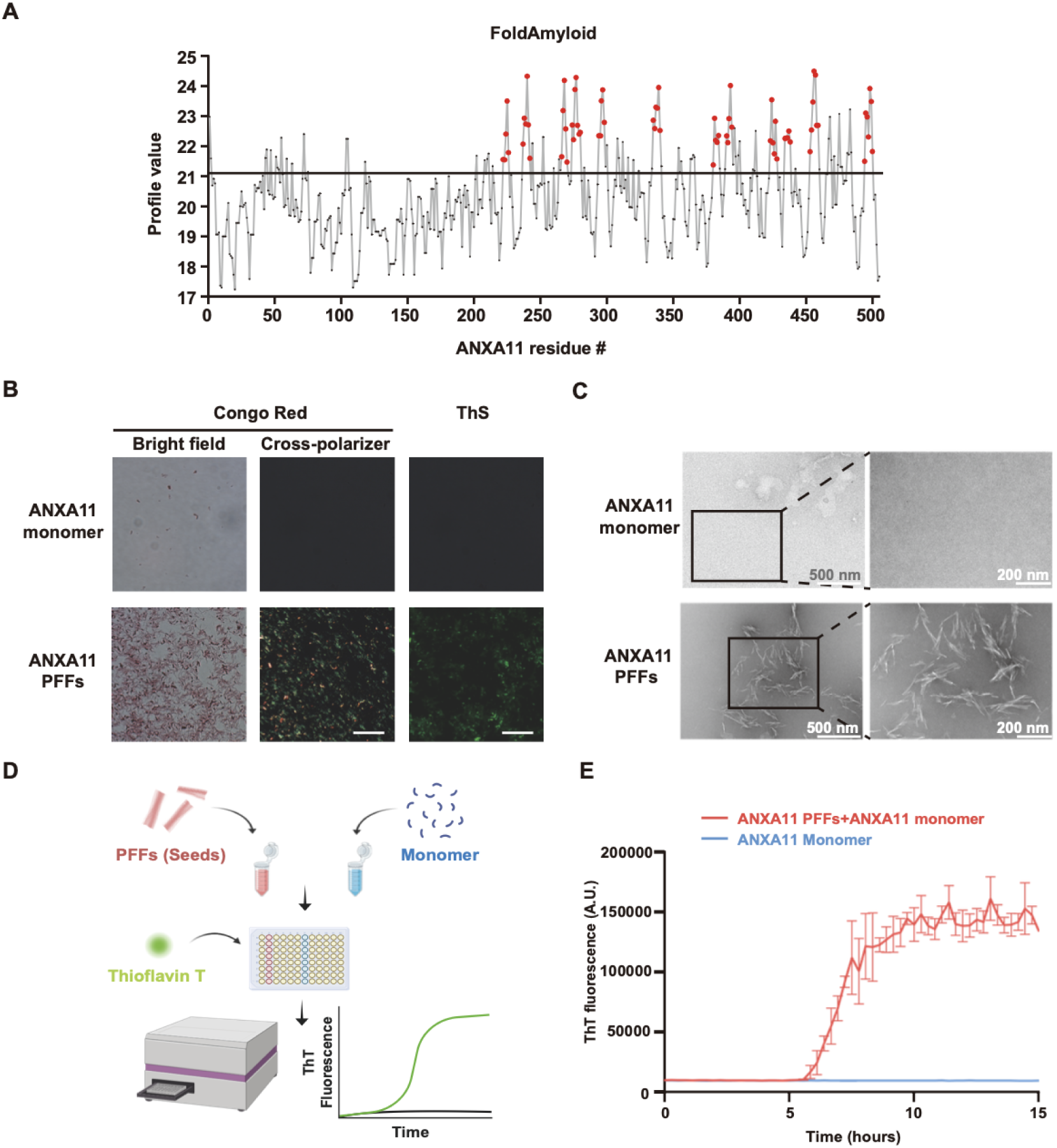
ANXA11 forms amyloid fibrils with prion-like self-templating activity. (A) FoldAmyloid prediction identifying aggregation-prone regions within the ANXA11 sequence. Regions above the threshold (red dashed line) indicate high amyloidogenic propensity. (B) Congo red and Thioflavin S (ThS) staining of aged ANXA11 assemblies. Congo red birefringence under polarized light (left) and ThS fluorescence (right) confirm amyloid nature. Scale bar: 50 μm. (C) Transmission electron microscopy (TEM) images comparing monomeric ANXA11 (top) with aged fibrillar ANXA11 (bottom). Fibrils display characteristic unbranched morphology. Scale bar: 100 nm. (D) Schematic of real-time quaking-induced conversion (RT-QuIC) assay for detecting prion-like seeding activity. Sonicated ANXA11 preformed fibrils (PFFs) are used to seed aggregation of monomeric ANXA11 substrate. (E) RT-QuIC kinetics showing seeding activity of ANXA11 PFFs. Thioflavin T (ThT) fluorescence intensity is plotted against time. Seeded reactions (colored lines) show accelerated aggregation compared to unseeded controls (black). Data represent mean ± SD from 3 independent experiments performed in triplicate.

Prion-like proteins can template the conversion of their native counterparts—a property that underlies the progressive nature of neurodegenerative diseases. To assess whether ANXA11 fibrils possess such self-templating activity, we employed real-time quaking-induced conversion (RT-QuIC), a highly sensitive assay for detecting prion-like seeding (Figure 3D). Sonicated ANXA11 preformed fibrils (PFFs) efficiently seeded the aggregation of monomeric ANXA11 substrate, as evidenced by a dramatic acceleration of ThT fluorescence increase compared to unseeded controls (Figure 3E). The seeding reaction eliminated the characteristic lag phase of spontaneous fibrillization. These data establish that ANXA11 fibrils exhibit hallmark prion-like properties, including self-templating seeding activity.

### ANXA11 fibrils propagate between cells and human iPSC-derived neurons

The prion-like spread of protein aggregates between cells is thought to underlie the progressive nature of neurodegenerative diseases. To assess whether ANXA11 aggregates can propagate between cells, we employed a transmission assay in SH-SY5Y neuroblastoma cells (Figure 4A). Cells were exposed to two different fluorescently labeled ANXA11 PFFs (Alexa Fluor 488 or Alexa Fluor 647) separately and then co-cultured to assess intercellular transmission. Confocal microscopy revealed uptake and transmission of ANXA11 aggregates between cells (Figure 4B). Flow cytometry-based quantification demonstrated efficient internalization and cell-to-cell transfer of ANXA11 aggregates (Figure 4C). Imaging flow cytometry further confirmed the internalization and transmission of ANXA11 aggregates between cells (Figure 4D).

**Figure 4.**
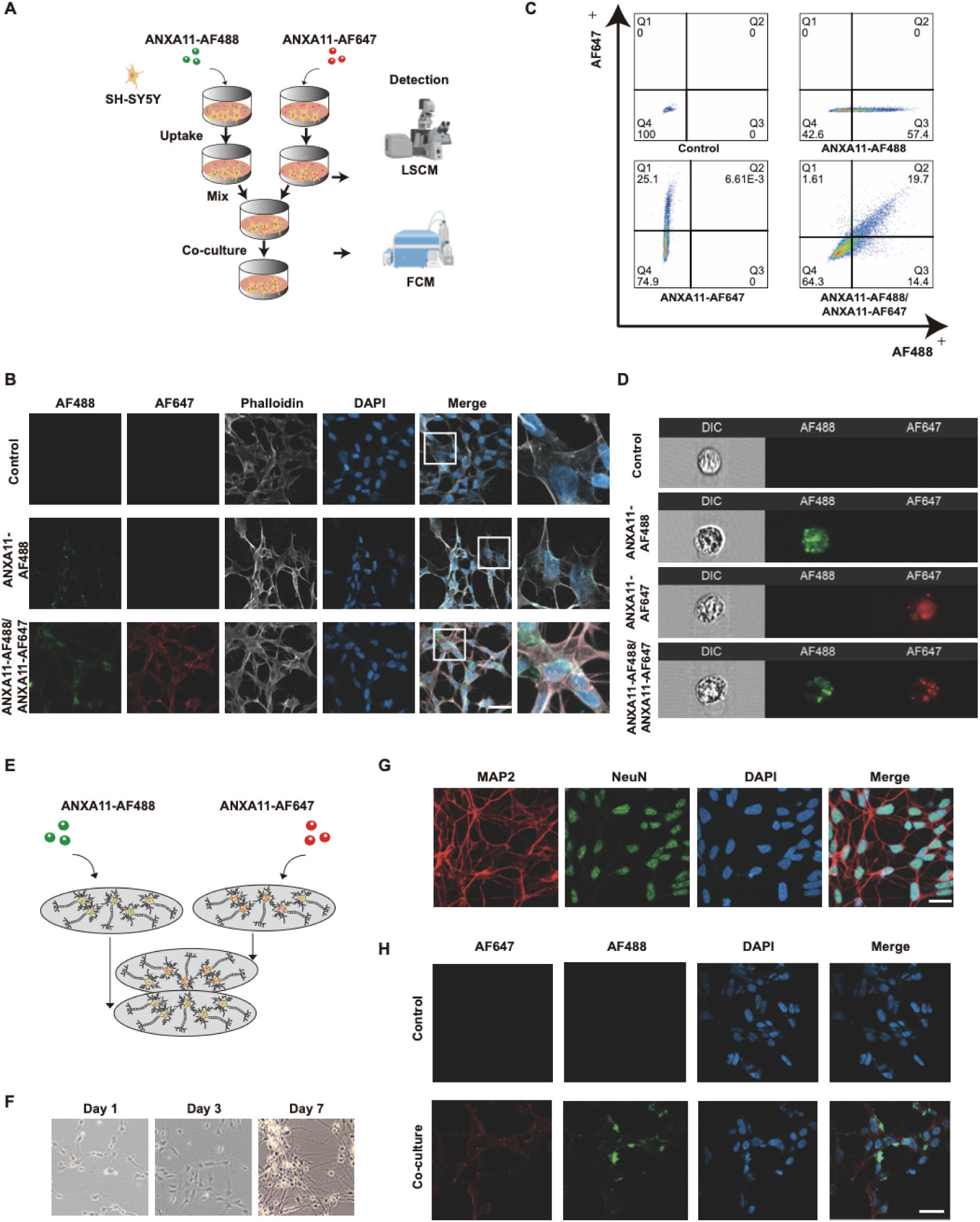
ANXA11 fibrils propagate between cells and human neurons. (A) Schematic of ANXA11 uptake and transmission assay in SH-SY5Y cells. Cells were exposed to two different fluorescently labeled ANXA11 PFFs separately and then co-cultured to assess intercellular transmission. (B) Confocal imaging of ANXA11 PFF uptake and transmission in SH-SY5Y cells. Alexa Fluor 488-labeled (green) and Alexa Fluor 647-labeled (red) PFFs are shown. Phalloidin (gray, F-actin), DAPI (blue, nuclei). Scale bar: 20 μm. (C) Flow cytometry-based quantification of ANXA11 transmission between SH-SY5Y cells. Percentage of recipient cells containing Alexa Fluor-labeled transferred ANXA11 is shown. Data are representative of 3 independent experiments. (D) Imaging flow cytometry confirming internalization and transmission of ANXA11 aggregates between cells. Representative images of individual cells are shown. (E) Schematic of iPSC-derived neuron transmission assay. Differentiated neurons were treated with AF488-labeled or AF647-labeled ANXA11 PFFs separately and then co-cultured to assess intercellular transmission. (F) Phase-contrast imaging of iPSC differentiation into cortical neurons showing characteristic neuronal morphology. Scale bar: 100 μm. (G) Immunofluorescence staining confirming neuronal identity of iPSC-derived neurons. MAP2 (green), NeuN (red), DAPI (blue). Scale bar: 50 μm. (H) Confocal imaging of neuron-to-neuron transmission of ANXA11 aggregates in iPSC-derived neurons. White arrowheads indicate transferred aggregates in recipient neurons. Scale bar: 20 μm.

To assess the disease relevance of ANXA11 propagation, we examined whether ANXA11 fibrils can spread between human neurons. We also employed a transmission assay in cortical neurons differentiated from human induced pluripotent stem cells (iPSCs) using established protocols (Figure 4E). Phase-contrast imaging confirmed successful neuronal differentiation with characteristic neuronal morphology (Figure 4F), and immunostaining for MAP2 and NeuN verified neuronal identity (Figure 4G). Upon treatment with two Alexa Fluor 488- or Alexa Fluor 647-labeled ANXA11 PFFs separately and then co-cultured, we observed intercellular transfer of ANXA11 aggregates between iPSC-derived neurons (Figure 4H). Together, these findings demonstrate that ANXA11 fibrils can propagate between human neurons, supporting the capacity of ANXA11 assemblies for intercellular transmission in a disease-relevant neuronal context.

### ANXA11 fibrils seed homotypic aggregation and induce TDP-43 pathological conversion

Having established that ANXA11 fibrils exhibit prion-like self-templating activity, we next examined their seeding capacity in cells. To test homotypic seeding, cells expressing ANXA11-GFP were treated with ANXA11 PFFs or vehicle control. Immunoblot analysis of detergent-soluble and detergent-insoluble fractions revealed that PFF treatment induced accumulation of ANXA11-GFP in the insoluble fraction (Figure 5A). Dot blot analysis with OC and A11 antibodies confirmed that PFF-seeded ANXA11 acquired amyloid-like properties, with increased reactivity with both antibodies (Figure 5B). Immunofluorescence microscopy of HEK293T cells demonstrated the formation of cytoplasmic ANXA11-GFP aggregates specifically in PFF-treated cells, with quantification confirming a significant increase in aggregate-positive cells (Figure 5C).

**Figure 5.**
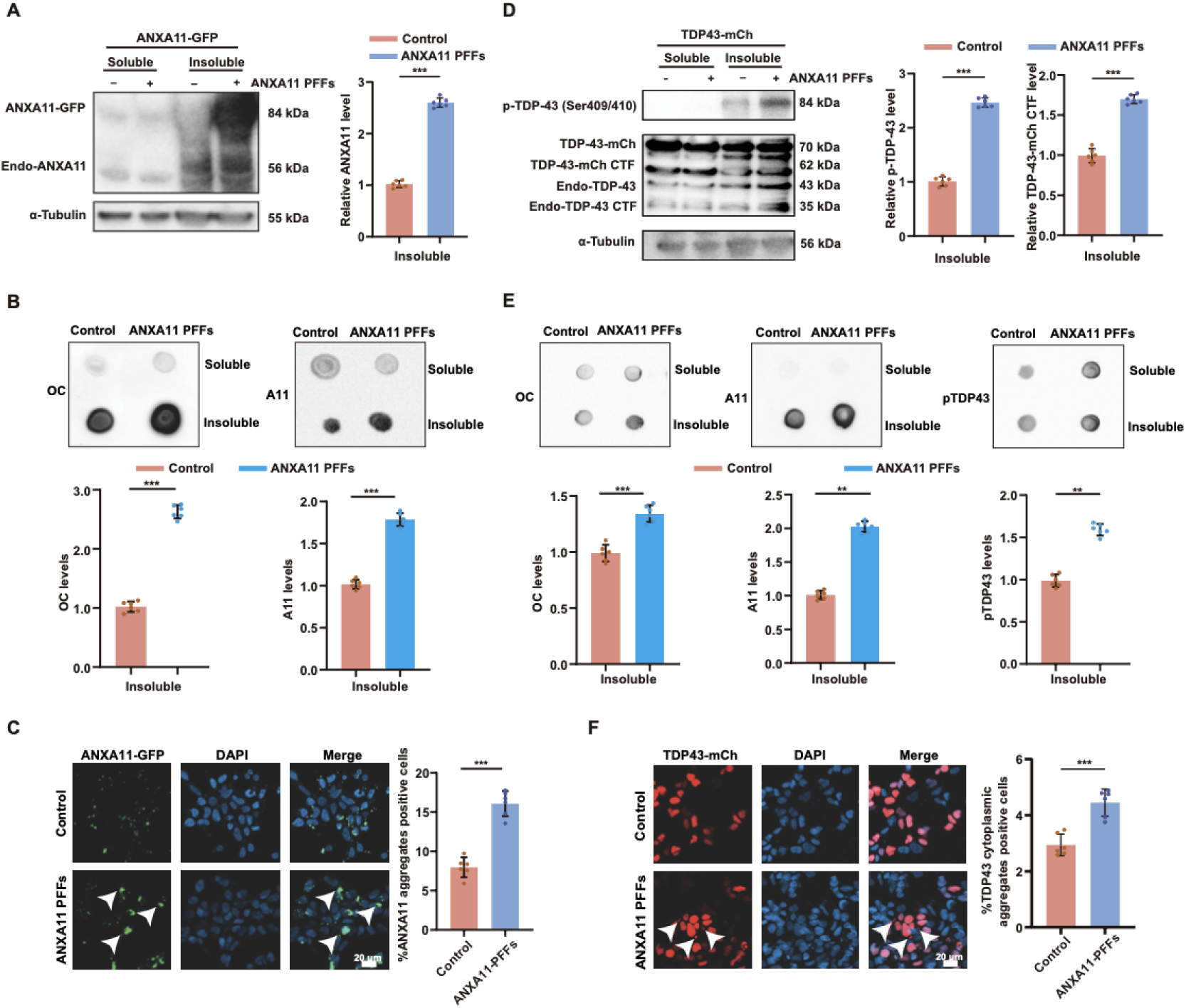
ANXA11 fibrils seed homotypic aggregation and cross-seed TDP-43 pathology. (A) Immunoblot analysis of detergent-soluble and -insoluble fractions from ANXA11-GFP-expressing cells treated with ANXA11 PFFs or vehicle control. PFF treatment induces accumulation of ANXA11-GFP in the insoluble fraction. α-Tubulin serves as loading control. (B) Dot blot analysis of ANXA11 seeded soluble and insoluble fractions from (A) using conformation-specific antibodies: OC (fibrillar amyloid) and A11 (prefibrillar oligomers). PFF-seeded ANXA11 shows increased reactivity with both antibodies. (C) Representative fluorescence images of ANXA11-GFP in HEK293T cells treated with vehicle or ANXA11 PFFs. ANXA11 PFFs treatment increased cytoplasmic puncta (ANXA11 aggregates; arrowheads). Quantification of aggregate-positive cells shown at right. Scale bar: 20 μm. (D) Immunoblot of detergent-soluble and detergent-insoluble fractions from HEK293T cells expressing TDP-43-mCherry treated with vehicle or ANXA11 PFFs. ANXA11 PFFs increased insoluble TDP-43-mCherry and phosphorylated TDP-43 (Ser409/410), and increased TDP-43 C-terminal fragments (CTF). Quantification shown at right (mean ± SEM; \*\*\**P* < 0.001). (E) Dot-blot analysis of soluble/insoluble fractions probed with OC, A11, and pTDP-43 (Ser409/410). ANXA11 PFFs treatment increased insoluble amyloid-reactive species and pTDP-43 signal. Quantification shown below (mean ± SEM; **P < 0.01, ***P < 0.001). (F) Representative images of TDP-43-mCherry-expressing cells treated with vehicle or ANXA11 PFFs. ANXA11 PFFs treatment increased cytoplasmic/extranuclear TDP-43 aggregates (arrowheads). Quantification shown at right. Scale bar, 20 μm.

The recent discovery of heteromeric ANXA11-TDP-43 filaments in FTLD-TDP type C raises the possibility that ANXA11 assemblies may promote TDP-43 pathological conversion. To test this, HEK293T cells expressing TDP-43-mCherry were exposed to ANXA11 PFFs. Immunoblot analysis showed that ANXA11 PFF treatment increased detergent-insoluble TDP-43-mCherry, enhanced phosphorylation of TDP-43 at Ser409/410, and promoted the appearance of C-terminal fragments (CTFs), all established hallmarks of pathological TDP-43 processing (Figure 5D). Dot blot analysis with OC, A11, and phospho-TDP-43 antibodies further confirmed increased accumulation of insoluble amyloid-reactive species and pTDP-43 signal following ANXA11 PFF treatment (Figure 5E). Immunofluorescence microscopy demonstrated the formation of cytoplasmic/extranuclear TDP-43-mCherry aggregates in ANXA11 PFF-treated cells, with quantification confirming a significant increase in aggregate-[ositive cells (Figure 5F). Together, these findings demonstrate that ANXA11 fibrils can induce pathological conversion of TDP-43 in cells, consistent with a heterotypic seeding mechanism and supporting a mechanistic link between ANXA11 aggregation and TDP-43 proteinopathy.

### TurboID proximity-labeling reveals enrichment of nuclear pore complex components in ANXA11 aggregate-proximal proteome

To identify cellular pathways engaged by ANXA11 aggregates, we employed TurboID proximity-labeling proteomics, which enables the identification of proteins in close proximity to a bait protein (Figure 6A). We generated stable cell lines expressing GFP-TurboID (control) or ANXA11-GFP-TurboID and induced aggregation by treatment with ANXA11 PFFs. Streptavidin immunoblotting of soluble and detergent-insoluble fractions confirmed robust proximity labeling with distinct fraction-dependent labeling patterns (Figure 6B). Biotinylated proteins were enriched by streptavidin pulldown and identified by mass spectrometry. Comparison of the ANXA11 interactome between soluble and insoluble fractions revealed aggregation-dependent rewiring, with distinct protein sets identified in each condition (Figure 6C).

**Figure 6.**
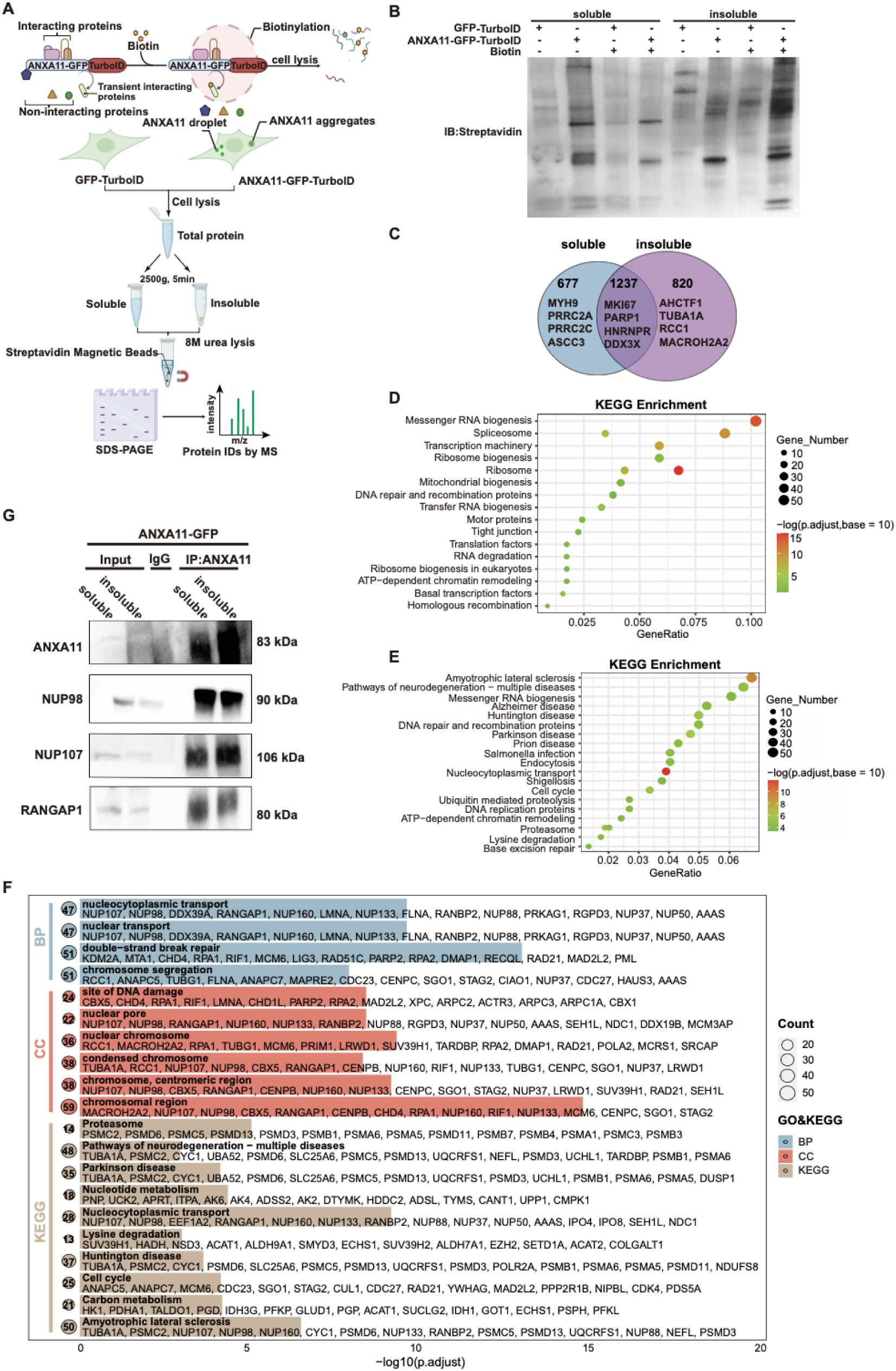
TurboID proximity labeling reveals enrichment of nuclear pore complex in ANXA11 aggregates. (A) Schematic of TurboID proximity-labeling proteomics workflow. Cells expressing ANXA11-TurboID are treated with ANXA11 PFFs or vehicle control. Biotinylated proteins are enriched by streptavidin pulldown and identified by mass spectrometry. (B) Streptavidin immunoblot of soluble and detergent-insoluble fractions from GFP-TurboID and ANXA11-GFP-TurboID cells (± biotin), showing robust proximity labeling and distinct fraction-dependent labeling patterns. (C) Venn diagram showing overlap of proteins identified in soluble versus insoluble ANXA11 proteomes. Numbers indicate proteins unique to each fraction or shared between fractions. (D) KEGG pathway enrichment analysis of the soluble ANXA11 proteome. Top enriched pathways include membrane trafficking and calcium signaling. Dot size represents gene count; color represents adjusted p-value. (E) KEGG pathway enrichment analysis of the insoluble ANXA11 proteome showing distinct pathway enrichment compared to soluble fraction. (F) Gene Ontology (GO) and KEGG enrichment analysis of proteins uniquely enriched in the insoluble fraction. Nuclear pore complex (NPC) components and nucleocytoplasmic transport machinery are significantly enriched. (G) Co-immunoprecipitation validation of ANXA11 interaction with NPC components. Cells expressing ANXA11-Flag were treated with ANXA11 PFFs or vehicle control. Lysates were immunoprecipitated with anti-Flag antibody and blotted for NUP98, NUP107, and RANGAP1. Input (5%) and IP fractions are shown.

KEGG pathway enrichment analysis of the soluble ANXA11 proteome revealed expected associations with membrane trafficking and calcium signaling (Figure 6D). In contrast, the insoluble ANXA11 proteome showed enrichment for a distinct set of pathways (Figure 6E). Strikingly, Gene Ontology and KEGG analyses of proteins uniquely enriched in the insoluble fraction revealed unexpected enrichment of nuclear pore complex (NPC) components and nucleocytoplasmic transport machinery among the most prominent categories (Figure 6F). These included multiple nucleoporins and transport-associated proteins, such as NUP98, NUP107, and the Ran GTPase-activating protein RANGAP1. Co-immunoprecipitation in cells expressing ANXA11-Flag treated with ANXA11 PFFs or vehicle control further confirmed aggregation-dependent association of ANXA11 with these factors (Figure 6G). Together, these findings indicate that ANXA11 aggregation is accompanied by marked rewiring of its proximal proteome, with prominent enrichment of NPC and nucleocytoplasmic transport components.

### ANXA11 aggregation disrupts NPC, impairs mRNA export, and causes neuronal toxicity

The enrichment of NPC components in the ANXA11 aggregate interactome prompted us to examine whether ANXA11 aggregation disrupts NPC integrity. Immunofluorescence microscopy of SH-SY5Y cells treated with buffer or ANXA11 PFF fibrils revealed pronounced nuclear envelope distortion and invaginations in aggregate-bearing cells. Cells harboring ANXA11 aggregates exhibited disruption of nuclear lamina organization, as evidenced by altered Lamin B1 distribution (Figure 7A). NPC components also showed striking redistribution in aggregate-bearing cells: NUP98 exhibited reduced nuclear rim staining and increased cytoplasmic puncta that co-localized with ANXA11 aggregates (Figure 7B), NUP107 lost its characteristic nuclear envelope pattern (Figure 7C), and RANGAP1 showed abnormal redistribution (Figure 7D). Quantification confirmed a significant increase in the propotion of cells with disrupted NPC-associated staining pattern following ANXA11 PFF treatmented (Figure 7E). Transmission electron microscopy further revealed ultrastructural abnormalities of the nuclear envelope in cells containing ANXA11 aggregates (Figure 7F). Together, these data indicate thatANXA11 aggregation induces marked remodeling of nuclear envelope and NPC architecture.

**Figure 7.**
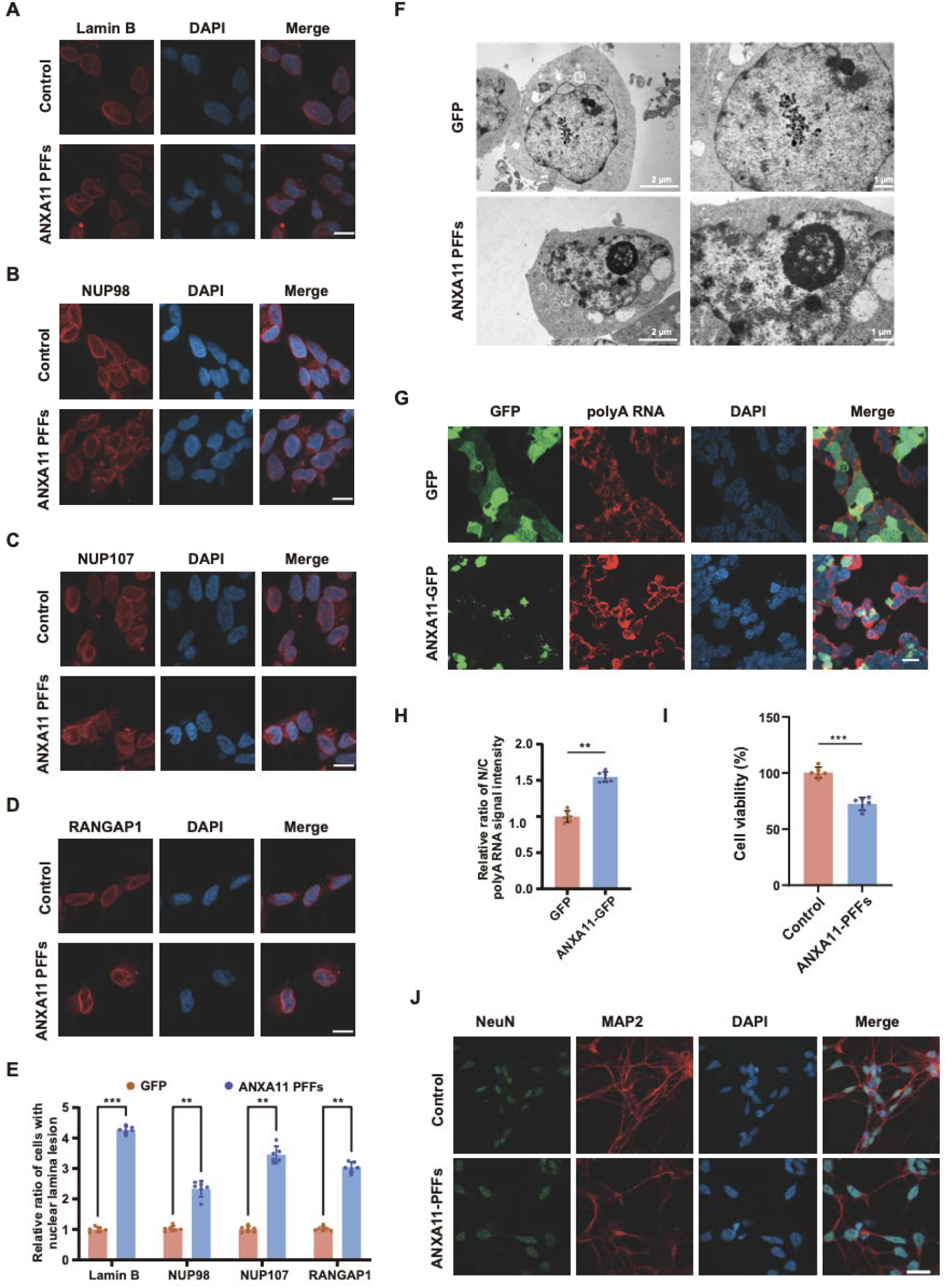
ANXA11 aggregation disrupts nuclear pore integrity, impairs mRNA export, and induces neuronal toxicity. (A-D) Immunofluorescence staining of Lamin B1 (A), NUP98 (B), NUP107 (C), and RANGAP1 (D) in SH-SY5Y cells treated with buffer (Control) or ANXA11 PFF fibrils. ANXA11 fibrils induce pronounced nuclear envelope distortion and invaginations. DAPI (blue) marks nuclei. Scale bars, 10 μm. Scale bar: 20 μm. (E) Quantification of NPC abnormalities. Percentage of cells with disrupted NPC staining pattern is shown for each marker. Data represent mean ± SD from at least 100 cells per condition from 3 independent experiments. Unpaired t-test; ****p < 0.0001. (F) Transmission electron microscopy (TEM) showing nuclear envelope morphology in control cells and cells with ANXA11 aggregates. Note ultrastructural abnormalities at the nuclear envelope in aggregate-bearing cells. NE, nuclear envelope; Cyt, cytoplasm; Nuc, nucleus. Scale bar: 500 nm. (G) Fluorescence in situ hybridization (FISH) using oligo-dT probes to detect polyadenylated mRNA. Poly(A)+ RNA (magenta), ANXA11 (green), DAPI (blue). Note nuclear accumulation of poly(A)+ RNA in cells with ANXA11 aggregates, indicating impaired mRNA export. Scale bar: 20 μm. (H) Quantification of nuclear-to-cytoplasmic (N/C) ratio of poly(A)+ mRNA signal in control cells versus cells with ANXA11 aggregates. Data are from at least 50 cells per condition and representative of 3 independent experiments. Unpaired t-test; ****p < 0.0001. (I) Cell viability assay of iPSC-derived neurons treated with ANXA11 PFFs or vehicle control over time. ANXA11 PFF treatment induces progressive neuronal toxicity. Data are representative of 3 independent experiments. Two-way ANOVA with Sidak’s post hoc test; *p < 0.05; **p < 0.01; ***p < 0.001. (J) Representative immunofluorescence images of iPSC-derived neurons treated with vehicle or ANXA11 PFFs and stained for NeuN (green) and MAP2 (red). ANXA11 PFFs treatment is associated with reduced neurite integrity and decreased MAP2 signal. DAPI (blue) marks nuclei. Scale bar, 20 μm.

Given that NPC integrity is essential for nucleocytoplasmic transport, we next assessed whether ANXA11 aggregation affects mRNA export. Fluorescence in situ hybridization (FISH) using oligo-dT probes to detect polyadenylated mRNA revealed nuclear accumulation of poly(A)+ RNA in cells harboring ANXA11 aggregates (Figure 7G), indicating impaired mRNA export. Quantification of the nuclear-to-cytoplasmic (N/C) ratio of poly(A)+ mRNA signal confirmed a significant increase in aggregation-bearing cells compared to control cells (Figure 7H). This redistribution was accompanied by reduced cytoplasmic mRNA signal, suggesting a functional defect in nucleocytoplasmic mRNA trasnport.

Finally, we assessed whether ANXA11 aggregation causes neuronal toxicity, the defining feature of neurodegenerative disease. Human iPSC-derived neurons were treated with ANXA11 PFFs or vehicle control and monitored over time. Cell viability assays showed progressive loss of neuronal viability following ANXA11 PFF exposure (Figure 7I). Immunofluorescence staining for NeuN and MAP2 further revealed reduced neurite integrity and decreased MAP2 signal in PFF-treatment neurons (Figure 7J). Together, these findings establish that ANXA11 aggregation disrupts NPC integrity, impairs nucleocytoplasmic mRNA transport, and ultimately causes neuronal toxicity.

## Discussion

Here, we demonstrate that ANXA11 possesses intrinsic amyloidogenic properties, undergoing liquid-liquid phase separation followed by a liquid-to-solid phase transition to amyloid fibrils with prion-like characteristics. We further show that ANXA11 fibrils exhibit self-templating seeding activity, propagate between cells, including iPSC-derived neurons, and induce pathological conversion of TDP-43 in cell models. In addition, TurboID proximity-labeling proteomics and cell-based analyses identify a strong association between ANXA11 aggregation and nucleocytoplasmic transport defects, including altered NPC-associated protein distribution and impaired poly(A)+ mRNA export. Together, these findings establish ANXA11 as a phase-transition-competent amyloidogenic protein and provide a mechanistic framework linking ANXA11 aggregation to TDP-43 co-pathology, transport dysfunction, and neuronal injury.

Our central conceptual advance is the demonstration that ANXA11 connects a physiologically plausible condensate state to a pathological amyloid state. The N-terminal intrinsically disordered region provides a structural basis for condensate formation (Smith et al., 2017; Liao et al., 2019), and the progressive loss of droplet dynamics observed in our FRAP analyses is consistent with emerging paradigms in which aging or cellular stress promotes condensation that hardens over time, increasing the likelihood of aberrant assembly (Mathieu et al., 2020; Visser et al., 2024). In this view, LLPS is not inherently pathological; rather, it creates concentrated microenvironments that can lower nucleation barriers and sensitize proteins to stress-, time-, or context-dependent conversion into stable β-sheet-rich assemblies (Lin et al., 2015; Molliex et al., 2015). These findings are particularly relevant to ANXA11 given its well-established roles at the interface of RNA granules and neuronal transport pathways (Liao et al., 2019; Nixon-Abell et al., 2025). The liquid-to-solid phase transition we observe in vitro was recapitulated in cellular models, where ANXA11-GFP condensates progressively acquired amyloid-like properties over time, as evidenced by OC and A11 antibody reactivity. This maturation process—from dynamic liquid droplets to solid-like aggregates—mirrors the pathological trajectory proposed for other neurodegenerative disease proteins including TDP-43 (Gasset-Rosa et al., 2019; Rummens et al., 2025) and FUS (Patel et al., 2015).

Our findings also support the view that ANXA11 shares key features with other aggregation-prone proteins, including TDP-43, tau, and α-synuclein, that undergo seeded amplification and intercellular transmission (Jucker and Walker, 2013; Lewis and Dickson, 2016; Nonaka et al., 2013). The RT-QuIC results provide strong evidence for self-templating conversion of ANXA11, demonstrating that preformed fibrils can accelerate aggregation of monomeric substrate without characteristic lag phase. This seeding activity is a hallmark of prion-like behavior and suggests that ANXA11 amyloid assemblies can catalyze further conversion of soluble protein, potentially amplifying proteopathic burden over time. In cell-based assays, ANXA11 fibrils were efficiently internalized and transmitted between SH-SY5Y cells and iPSC-derived neurons. Although these models do not define dissemination routes in vivo, they establish that ANXA11 amyloids can persist within cells and retain seeding activities after uptake, properties expected to drive progressive pathology. The propagation mechanism may partly explain the multisystem involvement observed in patients with ANXA11-related neurodegeneration (Leoni et al., 2021; Liu et al., 2025).

An important disease-relevant implication of our findings is that ANXA11 fibrils can induce pathological conversion of TDP-43. Exposure to ANXA11 fibrils increased detergent-insoluble TDP-43, phosphorylation at Ser409/410, C-terminal fragmentation, and the formation of cytosolic aggregates. These findings are consistent with a heterotypic seeding model and provide a plausible mechanism by which ANXA11 pathology can potentiate canonical TDP-43 proteinopathy, consistent with the recent identification of heteromeric ANXA11-TDP-43 amyloid filaments in FTLD-TDP type C patient brains (Arseni et al., 2024). In this context, ANXA11 aggregation may represent an upstream or parallel proteopathic event capable of promoting canonical TDP-43 pathology and amplifying disease-associated proteinopathy through a feed-forward mechanism.

Our proteomic and cellular analyses further converge on nucleocytoplasmic transport dysfunction as a major consequence of ANXA11 aggregation. Because mature ANXA11 assemblies are difficult to analyze using conventional extraction-based approaches, TurboID proximity labeling provided a useful strategy to define aggregation-associated changes in the ANXA11 proximal proteome (Branon et al., 2018). Strikingly, the insoluble ANXA11 proximity proteome was enriched for nuclear pore complex components and nucleocytoplasmic transport factors, including Nup98, Nup107, and RanGAP1. Nucleocytoplasmic transport mediates bidirectional macromolecule exchange across the nuclear envelope and depends on NPC integrity and nucleoporin barrier functions (Cook et al., 2007). Defects in this pathway are increasingly connected to neurodegenerative disease, particularly ALS/FTD (Kim and Taylor, 2017; Chou et al., 2018). Consistent with this, ANXA11 fibril exposure produced nuclear envelope distortions, altered NPC-associated staining, and lamin B mislocalization—resembling defects caused by disease-related ANXA11 mutations (Marchica et al., 2025). Functionally, ANXA11 fibrils impaired mRNA export, leading to nuclear accumulation of poly(A)+ RNA. This connects aggregation-dependent NPC disruption to defective nucleocytoplasmic output, providing a mechanistic link between ANXA11 pathology and cellular dysfunction. The convergence of ANXA11 and TDP-43 pathologies on nuclear transport dysfunction is particularly noteworthy, as TDP-43 aggregation has also been shown to disrupt NPC function (Chou et al., 2018), suggesting that these proteins may act synergistically to impair this critical cellular process. Ultimately, ANXA11 fibril exposure induced progressive toxicity in iPSC-derived human neurons, establishing the disease relevance of our findings.

Several limitations of our study warrant consideration. First, while our cellular and neuronal models recapitulate important features of ANXA11 aggregation, they do not fully capture the complexity of human disease, including the contribution of aging, genetic modifiers, and cell-type-specific vulnerabilities. Second, our seeded model is based primarily on full-length ANXA11 assemblies and therefore does not yet determine whether the active pathogenic species or interaction interfaces fully correspond to those present in patient-derived ANXA11-TDP-43 filaments. Third, while our data support ANXA11-induced pathological conversion of TDP-43, they do not yet establish the precise structural basis of heterotypic seeding or define the extent of endogenous nuclear TDP-43 loss of function. Fourth, proximity labeling and co-immunoprecipitation indicate aggregation-associated engagement of NPC factors, but do not by themselves distinguish direct molecular sequestration from indirect remodeling of the surrounding cellular environment. Future studies should address whether ANXA11 pathology precedes or follows TDP-43 aggregation in patient tissues, and whether targeting ANXA11 phase transition or propagation can prevent downstream TDP-43 pathology.

In conclusion, our study identifies ANXA11 as an intrinsically amyloidogenic, phase-transition-competent protein whose seeded assemblies propagate between cells, induce TDP-43 co-pathology, and are associated with nucleocytoplasmic transport defects. By linking ANXA11 phase transition to self-templating assembly, neuronal propagation, transport impairment, and neuronal injury, this work provides a mechanistic framework for understanding how ANXA11 aggregation may contribute to ALS/FTD pathogenesis. These findings also highlight ANXA11 phase behavior, seeded assembly, and nucleocytoplasmic transport vulnerability as candidate therapeutic entry points in ANXA11-associated neurodegeneration.

## Supplementary Figures

**Supplementary Figure 1:**
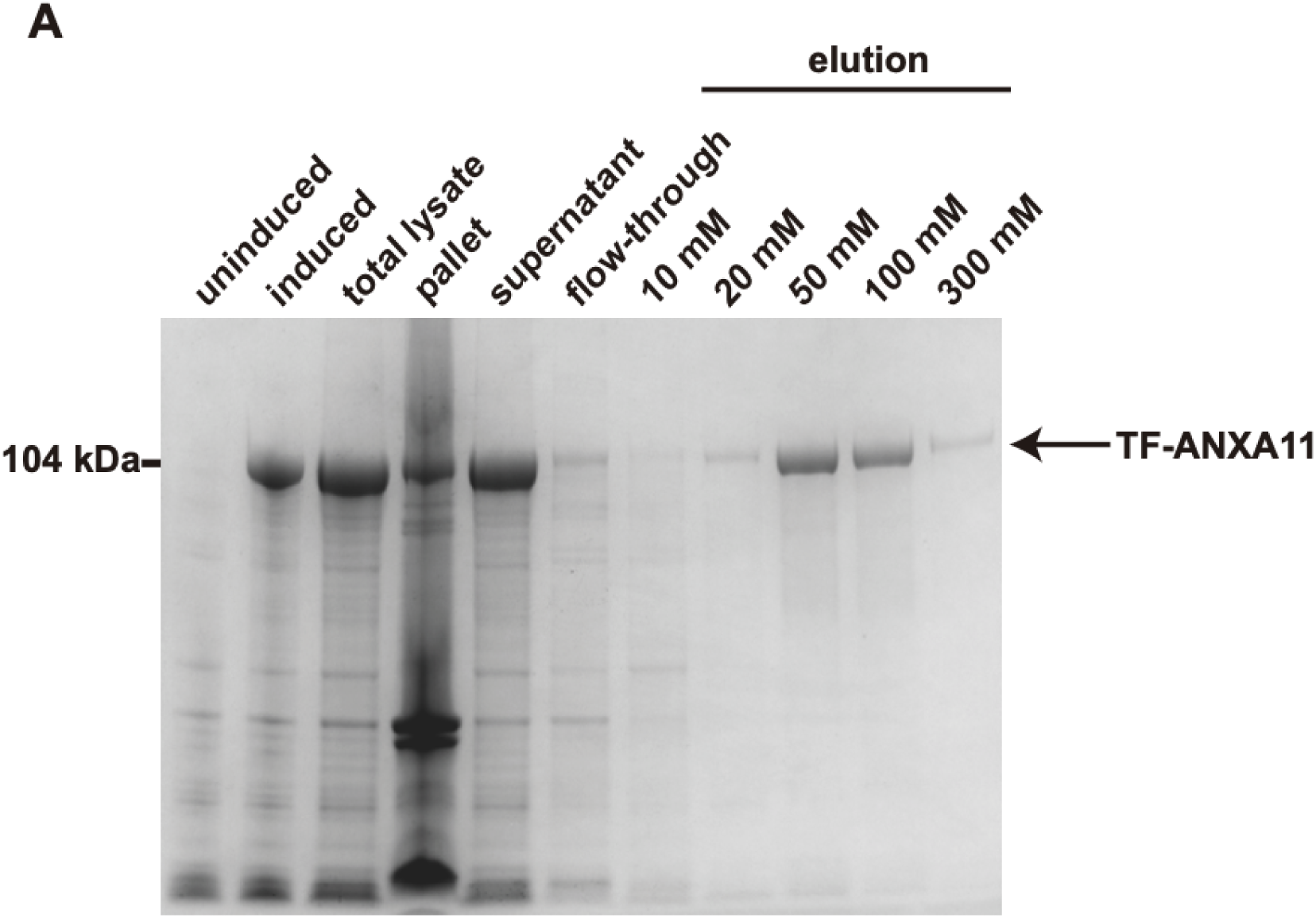
representation of purification of recombinant ANXA11.

## REFERENCES

1. Arai, T., Hasegawa, M., Akiyama, H., Ikeda, K., Nonaka, T., Mori, H., Mann, D., Tsuchiya, K., Yoshida, M., Hashizume, Y., and Oda, T. (2006). TDP-43 is a component of ubiquitin-positive tau-negative inclusions in frontotemporal lobar degeneration and amyotrophic lateral sclerosis. Biochem. Biophys. Res. Commun. 351, 602–611. DOI: 10.1016/j.bbrc.2006.10.093 PubMed: https://pubmed.ncbi.nlm.nih.gov/17084815/

2. Arseni, D., Nonaka, T., Jacobsen, M.H., Murzin, A.G., Cracco, L., Peak-Chew, S.Y., Garringer, H.J., Kawakami, I., Suzuki, H., Onaya, M., et al. (2024). Heteromeric amyloid filaments of ANXA11 and TDP-43 in FTLD-TDP type C. Nature 634, 662–668. DOI: 10.1038/s41586-024-08024-5 PubMed: https://pubmed.ncbi.nlm.nih.gov/39358506/

3. Branon, T.C., Bosch, J.A., Sanchez, A.D., Udeshi, N.D., Svinkina, T., Carr, S.A., Feldman, J.L., Perrimon, N., and Ting, A.Y. (2018). Efficient proximity labeling in living cells and organisms with TurboID. Nat. Biotechnol. 36, 880–887. DOI: 10.1038/nbt.4201 PubMed: https://pubmed.ncbi.nlm.nih.gov/30125270/

4. Chou, C.C., Zhang, Y., Umoh, M.E., Vaughan, S.W., Lorenzini, I., Liu, F., Sayegh, M., Donlin-Asp, P.G., Chen, Y.H., Duong, D.M., et al. (2018). TDP-43 pathology disrupts nuclear pore complexes and nucleocytoplasmic transport in ALS/FTD. Nat. Neurosci. 21, 228–239. DOI: 10.1038/s41593-017-0047-3 PubMed: https://pubmed.ncbi.nlm.nih.gov/29311743/

5. Cook, A., Bono, F., Jinek, M., and Conti, E. (2007). Structural biology of nucleocytoplasmic transport. Annu. Rev. Biochem. 76, 647–671. DOI: 10.1146/annurev.biochem.76.052705.161529 PubMed: https://pubmed.ncbi.nlm.nih.gov/17328675/

6. Gasset-Rosa, F., Lu, S., Yu, H., Chen, C., Melamed, Z., Guo, L., Shorter, J., Da Cruz, S., and Cleveland, D.W. (2019). Cytoplasmic TDP-43 de-mixing independent of stress granules drives inhibition of nuclear import, loss of nuclear TDP-43, and cell death. Neuron 102, 339–357.e7. DOI: 10.1016/j.neuron.2019.02.038 PubMed: https://pubmed.ncbi.nlm.nih.gov/30853299/

7. Jiang, Q., Lin, J., Wei, Q., Li, C., Hou, Y., Cao, B., Zhang, L., Ou, R., Liu, K., Yang, T., et al. (2022). Genetic analysis of and clinical characteristics associated with ANXA11 variants in a Chinese cohort with amyotrophic lateral sclerosis. Neurobiol. Dis. 175, 105907. DOI: 10.1016/j.nbd.2022.105907 PubMed: https://pubmed.ncbi.nlm.nih.gov/36280114/

8. Jucker, M., and Walker, L.C. (2013). Self-propagation of pathogenic protein aggregates in neurodegenerative diseases. Nature 501, 45–51. DOI: 10.1038/nature12481 PubMed: https://pubmed.ncbi.nlm.nih.gov/24005412/

9. Kim, H.J., and Taylor, J.P. (2017). Lost in transportation: nucleocytoplasmic transport defects in ALS and other neurodegenerative diseases. Neuron 96, 285–297. DOI: 10.1016/j.neuron.2017.07.029 PubMed: https://pubmed.ncbi.nlm.nih.gov/29024655/

10. Leoni, T.B., González-Salazar, C., Rezende, T.J.R., Hernández, A.L.C., Mattos, A.H.B., Coimbra Neto, A.R., da Graça, F.F., Gonçalves, J.P.N., Martinez, A.R.M., Taniguti, L., et al. (2021). A novel multisystem proteinopathy caused by a missense ANXA11 variant. Ann. Neurol. 90, 239–252. DOI: 10.1002/ana.26136 PubMed: https://pubmed.ncbi.nlm.nih.gov/34036626/

11. Lewis, J., and Dickson, D.W. (2016). Propagation of tau pathology: hypotheses, discoveries, and yet unresolved questions from experimental and human brain studies. Acta Neuropathol. 131, 27–48. DOI: 10.1007/s00401-015-1507-z PubMed: https://pubmed.ncbi.nlm.nih.gov/26576562/

12. Liao, Y.C., Fernandopulle, M.S., Wang, G., Choi, H., Hao, L., Drerup, C.M., Patel, R., Qamar, S., Nixon-Abell, J., Shen, Y., et al. (2019). RNA granules hitchhike on lysosomes for long-distance transport, using annexin A11 as a molecular tether. Cell 179, 147–164.e20. DOI: 10.1016/j.cell.2019.08.050 PubMed: https://pubmed.ncbi.nlm.nih.gov/31539493/

13. Lin, Y., Protter, D.S., Rosen, M.K., and Parker, R. (2015). Formation and maturation of phase-separated liquid droplets by RNA-binding proteins. Mol. Cell 60, 208–219. DOI: 10.1016/j.molcel.2015.08.018 PubMed: https://pubmed.ncbi.nlm.nih.gov/26412307/

14. Ling, S.C., Polymenidou, M., and Cleveland, D.W. (2013). Converging mechanisms in ALS and FTD: disrupted RNA and protein homeostasis. Neuron 79, 416–438. DOI: 10.1016/j.neuron.2013.07.033 PubMed: https://pubmed.ncbi.nlm.nih.gov/23931993/

15. Liu, Q., Sun, Y., He, B., Chen, H., Wang, L., Wang, G., Zhang, K., Zhao, X., Zhang, X., Shen, D., et al. (2025). Gain-of-function ANXA11 mutation cause late-onset ALS with aberrant protein aggregation, neuroinflammation and autophagy impairment. Acta Neuropathol. Commun. 13, 2. DOI: 10.1186/s40478-024-01915-0 PubMed: https://pubmed.ncbi.nlm.nih.gov/39754212/

16. Marchica, V., Biasetti, L., Barnard, J., Li, S., Nikolaou, N., Frosch, M.P., Lucente, D.E., Eldaief, M., King, A., Fanto, M., et al. (2025). Annexin A11 mutations are associated with nuclear envelope dysfunction in vivo and in human tissues. Brain 148, 276–290. DOI: 10.1093/brain/awae256 PubMed: https://pubmed.ncbi.nlm.nih.gov/39101559/

17. Mathieu, C., Pappu, R.V., and Taylor, J.P. (2020). Beyond aggregation: pathological phase transitions in neurodegenerative disease. Science 370, 56–60. DOI: 10.1126/science.abb8032 PubMed: https://pubmed.ncbi.nlm.nih.gov/33004511/

18. Molliex, A., Temirov, J., Lee, J., Coughlin, M., Kanagaraj, A.P., Kim, H.J., Mittag, T., and Taylor, J.P. (2015). Phase separation by low complexity domains promotes stress granule assembly and drives pathological fibrillization. Cell 163, 123–133. DOI: 10.1016/j.cell.2015.09.015 PubMed: https://pubmed.ncbi.nlm.nih.gov/26406374/

19. Murakami, T., Qamar, S., Lin, J.Q., Schierle, G.S., Bharat, T.A.M., Hainsworth, A.H., and St George-Hyslop, P. (2015). ALS/FTD mutation-induced phase transition of FUS liquid droplets and reversible hydrogels into irreversible hydrogels impairs RNP granule function. Neuron 88, 678–690. DOI: 10.1016/j.neuron.2015.10.030 PubMed: https://pubmed.ncbi.nlm.nih.gov/26526393/

20. Nahm, M., Lim, S.M., Kim, Y.E., Park, J., Noh, M.Y., Lee, S., Roh, J.E., Hwang, S.M., Park, C.K., Kim, Y.H., et al. (2020). ANXA11 mutations in ALS cause dysregulation of calcium homeostasis and stress granule dynamics. Sci. Transl. Med. 12, eaax3993. DOI: 10.1126/scitranslmed.aax3993 PubMed: https://pubmed.ncbi.nlm.nih.gov/32461334/

21. Neumann, M., Sampathu, D.M., Kwong, L.K., Truax, A.C., Micsenyi, M.C., Chou, T.T., Bruce, J., Schuck, T., Grossman, M., Clark, C.M., et al. (2006). Ubiquitinated TDP-43 in frontotemporal lobar degeneration and amyotrophic lateral sclerosis. Science 314, 130–133. DOI: 10.1126/science.1134108 PubMed: https://pubmed.ncbi.nlm.nih.gov/17023659/

22. Nixon-Abell, J., Ruggeri, F.S., Qamar, S., Herling, T.W., Czekalska, M.A., Shen, Y., Wang, G., King, C., Fernandopulle, M.S., Sneideris, T., et al. (2025). ANXA11 biomolecular condensates facilitate protein-lipid phase coupling on lysosomal membranes. Nat. Commun. 16, 2814. DOI: 10.1038/s41467-025-56636-8 PubMed: https://pubmed.ncbi.nlm.nih.gov/39929810/

23. Nonaka, T., Masuda-Suzukake, M., Arai, T., Hasegawa, Y., Akatsu, H., Obi, T., Yoshida, M., Murayama, S., Mann, D.M., Akiyama, H., and Hasegawa, M. (2013). Prion-like properties of pathological TDP-43 aggregates from diseased brains. Cell Rep. 4, 124–134. DOI: 10.1016/j.celrep.2013.06.007 PubMed: https://pubmed.ncbi.nlm.nih.gov/23831027/

24. Patel, A., Lee, H.O., Jawerth, L., Maharana, S., Jahnel, M., Hein, M.Y., Stoynov, S., Mahamid, J., Saha, S., Franzmann, T.M., et al. (2015). A liquid-to-solid phase transition of the ALS protein FUS accelerated by disease mutation. Cell 162, 1066–1077. DOI: 10.1016/j.cell.2015.07.047 PubMed: https://pubmed.ncbi.nlm.nih.gov/26317470/

25. Polymenidou, M., and Cleveland, D.W. (2011). The seeds of neurodegeneration: prion-like spreading in ALS. Cell 147, 498–508. DOI: 10.1016/j.cell.2011.10.011 PubMed: https://pubmed.ncbi.nlm.nih.gov/22036560/

27. Rummens, J., Khalil, B., Yıldırım, G., Silva, P., Zorzini, V., Peredo, N., Wojno, M., Ramakers, M., Van Den Bosch, L., Van Damme, P., et al. (2025). TDP-43 seeding induces cytoplasmic aggregation heterogeneity and nuclear loss of function of TDP-43. Neuron 113, 1597–1613. DOI: 10.1016/j.neuron.2025.03.004 PubMed: https://pubmed.ncbi.nlm.nih.gov/40154484/

28. Scialò, C., Zhong, W., Jagannath, S., Wilkins, O., Caredio, D., Hruska-Plochan, M., Lurati, F., Peter, M., De Cecco, E., Celauro, L., et al. (2025). Seeded aggregation of TDP-43 induces its loss of function and reveals early pathological signatures. Neuron 113, 1614–1628. DOI: 10.1016/j.neuron.2025.03.008 PubMed: https://pubmed.ncbi.nlm.nih.gov/40154485/

29. Smith, B.N., Topp, S.D., Fallini, C., Shibata, H., Chen, H.J., Troakes, C., King, A., Ticozzi, N., Kenna, K.P., Soragia-Gkazi, A., et al. (2017). Mutations in the vesicular trafficking protein annexin A11 are associated with amyotrophic lateral sclerosis. Sci. Transl. Med. 9, eaad9157. DOI: 10.1126/scitranslmed.aad9157 PubMed: https://pubmed.ncbi.nlm.nih.gov/28469040/

30. Sung, W., Bhardwaj, A., Bhardwaj, S., Bhardwaj, R., Bhardwaj, V., Bhardwaj, P., Bhardwaj, M., and Bhardwaj, K. (2024). ANXA11 mutations in amyotrophic lateral sclerosis: clinical, genetic, and functional characterization. Neurobiol. Aging 133, 1–10. DOI: 10.1016/j.neurobiolaging.2023.09.012 PubMed: https://pubmed.ncbi.nlm.nih.gov/37839265/

31. Vance, C., Rogelj, B., Hortobágyi, T., De Vos, K.J., Nishimura, A.L., Sreedharan, J., Hu, X., Smith, B., Ruddy, D., Wright, P., et al. (2009). Mutations in FUS, an RNA processing protein, cause familial amyotrophic lateral sclerosis type 6. Science 323, 1208–1211. DOI: 10.1126/science.1165942 PubMed: https://pubmed.ncbi.nlm.nih.gov/19251628/

32. Zhang, K., Donnelly, C.J., Haeusler, A.R., Grima, J.C., Machamer, J.B., Steinwald, P., Daley, E.L., Miller, S.J., Cunningham, K.M., Viber, S., et al. (2015). The C9orf72 repeat expansion disrupts nucleocytoplasmic transport. Nature 525, 56–61. DOI: 10.1038/nature14973 PubMed: https://pubmed.ncbi.nlm.nih.gov/26308891/

33. Zhang, Y.J., Gendron, T.F., Grima, J.C., Sasaguri, H., Jansen-West, K., Xu, Y.F., Katzman, R.B., Gass, J., Murray, M.E., Shinohara, M., et al. (2016). C9ORF72 poly(GA) aggregates sequester and impair HR23 and nucleocytoplasmic transport proteins. Nat. Neurosci. 19, 668–677. DOI: 10.1038/nn.4272 PubMed: https://pubmed.ncbi.nlm.nih.gov/26998601/

